## Supplementary Information for "An efficient urine peptidomics workflow identifies chemically defined dietary gluten peptides from patients with celiac disease"

**This PDF file includes:**

Supplementary Methods
Supplementary Figures 1 to 10
Supplementary Tables 1 to 3
Supplementary References

### SUPPLEMENTAL MATERIALS AND METHODS

#### Method Development

*Pilot Analysis of Urine from Individuals Challenged with Dietary Gluten.* In LC-MS/MS-based proteomic experiments, reversed-phase extraction techniques with C18 resins are commonly used to remove salts and small molecules from peptides prior to LC-MS/MS[1]. Therefore, we initially tested a typical C18 solid phase extraction (SPE) protocol for its suitability to urine samples for gluten-derived dietary peptide analysis.

Three volunteers were recruited to eat a meal consisting of two wheat bagels and to collect spot urine samples approximately 3-4 hours thereafter. These samples underwent initial processing, storage, reduction and alkylation as described the **Main Text Methods**. After these steps, urine (3-4.5 mL) was applied to 10 kDa cutoff centrifugal filtration units (Amicon Ultra-4, UFC800308) and centrifuged at 5000 x g until less than 100 µl of urine remained in the retentate. The filtrate was acidified with formic acid (FA) to a final concentration of 2% (v/v). The acidified filtrate was applied in 1 mL aliquots to C18 SPE columns (Agilent, #5982-1111) that had been pretreated with 1 mL of 100% acetonitrile and equilibrated with 2 times with 1-mL aliquots of 2% aqueous FA. Bound peptides were washed with 3 x 1 mL aliquots of 2% aqueous FA. Finally, peptides were eluted in 1 mL of 60% acetonitrile/38% water/2% FA directly into microcentrifuge tubes. For all SPE steps, a vacuum manifold was used to achieve a flow rate of ~1 drop per second. Optionally, in an attempt to remove urinary pigments and metabolites, 1 mL of ethyl acetate was added to the eluent, vortexed, and then centrifuged at 16,000 x g to separate the aqueous and organic layers. The ethyl acetate layer was discarded. The peptide-containing aqueous layer, which had a yellow color, was flash frozen on liquid nitrogen and dried by lyophilization.

At the time of LC-MS/MS analysis, samples were reconstituted in 25 µL MilliQ water. LC-MS/MS was performed essentially as previously reported in detail [2]. Briefly, 4 µL of reconstituted peptides were separated by capillary reverse phase chromatography on a 20 cm reversed-phase column packed in-house with C18 resin on an Eksigent Ekspert nanoLC-425 or a Dionex Ultimate 3000 system using a two-step linear gradient with 4–25% buffer B (0.1% v/v FA and 5% DMSO in acetonitrile) for 20 min followed by 25–40% buffer B for 5 min. MS/MS analysis was performed using an Orbitrap Elite mass spectrometer operated in the Top 20 data-dependent acquisition mode with collision-induced dissociation for peptide fragmentation. Data were searched as described in the **Main Text Methods**, with the exception that the fragment mass tolerance was set to 0.5 Da.

*Identification of a Sentinel Urine Sample for Method Optimization.* For optimizing our urine peptidomic cleanup protocol, we first sought to identify a gluten-positive urine specimen as a reference sample. To do so, a 12-hour pooled urine sample was collected from a dermatitis herpetiformis patient who reported long-term (>1 month) adherence to a GFD. On the day following this urine collection, the individual repeated the 12-hour urine collection after consuming two wheat bagels (~18 g gluten). Gliadin (one of two major protein families that make up gluten) content in these urine samples was confirmed by an R5 antibody-based competitive ELISA (R7021, R-Biopharm) following the manufacturer’s instructions.

*Optimization of Gluten Peptide Recovery by Centrifugal Filtration*. Prior reports of endogenous human urinary peptidome analysis by LC-MS/MS used centrifugal filtration to deplete high-molecular weight urinary proteins from lower-molecular weight peptides[3–6]. We first sought to verify that a similar strategy could enhance gluten peptide recovery while selecting the optimal molecular weight cutoff (MWCO). Centrifugal filters with three MWCOs were employed: 10 kDa, 30 kDa, and 50 kDa. After passing a urine sample through each filtration device, the retentate and flow-through were analyzed for gliadin peptide content by R5 ELISA, as described above, and for total protein content by SDS-PAGE and silver-stain densitometry.

*Optimization of Urinary Peptidomic Cleanup and LC-MS/MS Analysis Protocol.* To maximize the number of wheat-derived peptide identifications, we systematically varied the steps in our sample preparation and LC-MS/MS analysis method (**Supplementary Figure 3a**) using 4 mL of the gluten-positive reference urine processed using a 10 kDa MWCO filtration device as a reference (**Supplementary Figure 2**). We endeavored to minimize carryover of urinary salts and metabolites that could lead to instrument downtime. Notably, urochrome, the major urinary pigment, has a strong yellow color[7], and we were therefore able to visually inspect samples as a proxy for successful sample cleanup, as described below.

As a starting point, reference urine was processed using Method A (**Supplementary Figure 3**). This method is described in detail above (*Pilot Analysis of Urine from Individuals Challenged with Dietary Gluten*). After elution from the reversed-phase C18 solid phase extraction cartridge, extraction of urinary metabolites using ethyl acetate was attempted as suggested in the literature[3].

We hypothesized that some peptides remained in the retentate of the centrifugal filters due to non-specific binding of these peptides to urinary proteins. To disrupt these putative peptide-protein interactions, 400 µL acetonitrile (ACN) was added to urine (20% final ACN concentration) just before application to the centrifugal filter. This pre-filtration denaturation step tripled the number of wheat peptide identifications. However, it resulted in the need to remove acetonitrile by drying the sample on a lyophilizer before reconstitution in 2% FA to proceed with reversed-phase C18 SPE (Method B, **Supplementary Figure 3a**). This process added one day to the overall sample workup protocol. We tested whether the addition of concentrated FA to a final concentration of 2% prior to filtration would similarly be effective to denature protein-peptide interactions (Method C, **Supplementary Figure 3a**). Method C yielded approximately the same number of wheat peptide identifications and three times the number of human peptide identifications as Method B (**Supplementary Figure 3c**), while eliminating the need for an intermediate sample drying step steps. Therefore, pre-filtration acidification of urine was optimal for peptide identification. However, attempts to analyze successive urine samples prepared using Method C inevitably led to a decrease in instrument performance, presumably due to the presence of urochrome and other urinary metabolites that were not effectively removed by C18 SPE.

Recent urinary peptidomic analysis reports suggested offline HPLC fractionation by strong cation exchange (SCX) chromatography could be beneficial[5,6]. Although the rationale for this step was not explicitly stated, SCX could effectively separate urochrome, an acidic molecule, from basic peptides. Here, we sought to avoid the need for offline HPLC fractionation, as the added time would impede the throughput required for comparing large numbers of clinical samples. Instead, we sought to adapt a procedure for microscale peptide cleanup using SCX StageTips[8]. To this end, reference urine was processed by Method D, which included all steps in Method C followed by drying of the sample by lyophilization, resuspension in 100 µL 1% FA, and SCX StageTip processing (**Supplementary Figure 3a**).

To generate SCX StageTips, low-binding 200 µL pipet tips were packed with three layers of Empore SCX resin (3M, #2251) as previously described[8]. StageTips were conditioned with 100% ACN, then with 1 M NaCl, 50 mM $NH_4HCO_3$, and equilibrated three times with 1% FA in water. Resuspended peptide samples were applied and then washed once in 1% FA in water, then twice in 80% ACN, 18% water, and 2% FA. Peptides were eluted into low-binding microcentrifuge tubes with 500 mM $NH_4COOH$, 20% acetonitrile, and 0.4% FA. All steps were carried out using 100 µL of the indicated solution. Eluted peptides were dried under reduced pressure on a SpeedVac and then reconstituted in 25 µL MilliQ water. The SCX cleanup procedure appeared to effectively deplete most urochrome from the samples, as judged by a reduction in yellow color (**Supplementary Figure 3b**). Despite a slight decrease in the number of wheat peptide identifications relative to Method C (**Supplementary Figure 3c)**, we chose to include the SCX StageTip cleanup step in order to avoid LC-MS/MS downtime caused by urinary metabolite contamination.

Next, we optimized the LC/MS-MS acquisition method. Method D used a relatively short (25 min) LC gradient, as detailed in *Pilot Analysis of Urine from Individuals Challenged with Dietary Gluten*. An extended gradient of 90 min (Method E, **Supplementary Figure 3a**) nearly tripled the number of wheat peptide identifications (**Supplementary Figure 3c**). For all MS analyses of urine with Methods A-E, an Orbitrap Elite instrument was used, and MS2 fragments generated by collision-induced dissociation (CID) were analyzed at low resolution in the ion trap (fragment mass tolerance used for data searches = 0.5 Da). Next, we analyzed SCX-purified urinary peptides with a similar extended LC gradient, but using an Orbitrap Fusion instrument (Method F, **Supplementary Figure 3a**). Strikingly, analysis with Method F led to a 4-fold increase in wheat peptide identifications (**Supplementary Figure 3c**). The reasons for this increase were not pursued in detail, but we speculate a primary factor is that the increased scan speed of the Orbitrap Fusion allowed us to analyze CID-generated MS2 fragment ions at high resolution in the Orbitrap (fragment mass tolerance used for data searches = 0.02 Da). While Method F improved our overall ability to identify wheat peptides approximately 20-fold compared to Method A, Method F required two days to complete due to the need to dry samples between C18 SPE and SCX StageTip processing steps. Additionally, although SCX StageTip cleanup appeared to remove the majority of urochrome (**Supplementary Figure 3b**), we still noted occasional decreases in instrument performance after attempting to analyze large batches of samples (>10).

We hypothesized that MCX SPE columns (Waters Corporation #186008918) that contain a mixed-mode (reversed-phase and cation exchange) resin could replace separate C18 SPE and SCX StageTip extraction steps. Accordingly, Method G (carried out as described in the **Main Text Methods)** not only appeared to completely remove urochrome from urine samples (**Supplementary Figure 3b**), but it also resulted in a ~1.5 fold increase in the number of both wheat and human peptide identifications compared to Method F. Method G also required fewer sample processing steps than Methods A-F (**Supplementary Figure 3a**), allowing us to generate LC-MS/MS ready urinary peptide samples in under 6 hours. Thus, Method G was chosen as the final analysis protocol, with a minor modification that peptides with charge state 3 and higher were selected for EThcD fragmentation in addition to CID fragmentation as detailed in the **Main Text Methods**.

**Validation of Wheat-Derived Peptide Sequences Identified by PEAKS Software**

The output of the PEAKS software used to identify peptide sequence from the LC-MS/MS data suggested that some wheat peptides were present in their deamidated forms. The mass difference between a deamidated peptide and the second isotopic peak of a native peptide (containing one $^{13}C$) is only 0.0193 Da. Thus, aberrant deamidation assignments commonly occur in proteomics studies when the second isotopic peak of a native peptide is selected for fragmentation[9]. Given the physiological importance of deamidation in CeD pathogenesis, we sought to manually validate all wheat peptides where deamidation was automatically assigned by the PEAKS software, as described below.

The isotopic pattern of the MS1 (precursor ion) spectrum of each peptide assigned as deamidated by PEAKS was inspected in Thermo Xcalibur 3.0. To determine if the assignment was better explained by *bona fide* deamidation or selection of the second, or in a few cases, third isotopic peak, the theoretical mass spectra of both the native and deamidated peptides were calculated using the MS-Isotope module of ProteinProspector (http://prospector.ucsf.edu/). Then, the isotopic pattern and mass error of the experimentally observed spectra were compared to the theoretical mass spectra for both the native and deamidated forms of the peptide. If the monoisotopic mass of the native peptide was visible along with the non-monoisotopic mass that was selected for fragmentation, then the peptide sequence was manually reassigned from the deamidated to the native sequence. In cases where the monoisotopic mass of the native peptide was not visible, but the experimentally observed mass more closely matched the theoretical non-monoisotopic mass of the native peptide compared to the deamidated form, then the sequence was also manually reassigned as native. ~90% of the peptide sequences assigned as deamidated by PEAKS were corrected to their native sequences on the basis of these two criteria. In **Supplementary Datasets 1-7**, peptides whose PEAKS-assigned sequences were manually reassigned as native are clearly denoted in the “Manually Corrected Sequence” column.

For the remaining ~10% of peptides with putative deamidation assignments, the observed masses and isotopic patterns in the MS1 spectra were either equally consistent with the native and deamidated peptide forms, or more consistent with the deamidated form. In these cases, we inspected the MS2 (product ion) spectra for the presence of fragment ions that would allow us to assign the deamidation to a particular Gln residue within the peptide sequence. In all of these cases, we were unable to unambiguously identify a particular Gln residue that had undergone deamidation, as assessed by the presence of multiple site-determining fragment ions. Because the combined MS1- and MS2-level evidence these for these peptide sequences were not clearly more consistent with either the deamidated or the native peptide sequences, we did not consider these peptides for downstream data analysis. We cannot exclude the possibility that some peptides were indeed deamidated (or a mixture of native and deamidated forms) but produced insufficient quality spectra to regiospecifically assign the deamidation site. Alternatively, it is possible that the sequences of these peptides were otherwise misassigned by the PEAKS software. For clarity, all excluded peptides are reported in the PEAKS search output in **Supplementary Datasets 1-7** and are denoted “N” in the “Included for Data Analysis” column.

We acknowledge that this approach for validating deamidated peptide sequences is stringent. To assign a peptide as deamidated, we required that the isotopic pattern and mass errors of the MS1 spectra to be consistent with the deamidated form, and for the MS2 spectra to be of sufficient quality to allow clear assignment of the deamidation site. Future studies utilizing targeted mass spectrometry and synthetic peptide standards should be insightful in conclusively determining if deamidated wheat peptides are detectable in urine, and if so, how their concentration compares to the corresponding native forms.

**FASTA Database Construction for Identifying Wheat Derived Peptides**

The protein database used to search for peptides from the human and wheat proteomes (data in Figures 1, 3, and 4 of the Main Text), was curated from the UniProt repository (www.uniprot.com) as follows. Protein sequences from the SwissProt databases (containing manually curated protein sequences) of the human (*Homo sapiens*) and wheat (*Triticum aestivium*), taxonomies were concatenated into a single FASTA file. Additionally, all protein sequences from the wheat taxonomy that contained the terms “gliadin”, or “glutenin”, in the TrEMBL databases (containing automatically curated sequences from genome sequencing) were also downloaded and added to the FASTA file containing the SwissProt sequences. All sequences were downloaded from UniProt on December 7, 2019, and the FASTA file has been deposited in the PRIDE repository under accession number PXD023160.

**FASTA Database Construction for Identification of Wheat, Rye, and Barley Derived Peptides**

The protein database used to search for peptides from the human, wheat, barley, and rye proteomes, (data in **Figure 3** section of the Main Text), was curated from the UniProt repository as follows. Protein sequences from the SwissProt databases of the human (*Homo sapiens*), wheat (*Triticum aestivium*), rye (*Secale cereale*), and barley (*Hordeum vulgare*) taxonomies were concatenated into a single FASTA file. Additionally, all protein sequences from the wheat, rye, and barley taxonomies that contained the terms “gliadin”, “glutenin”, “secalin”, or “hordein” in the TrEMBL databases were also downloaded and added to the FASTA file containing the SwissProt sequences. All sequences were downloaded from UniProt on October 7, 2020, and the FASTA file has been deposited in the PRIDE repository under accession number PXD023160.

**Assignment of Peptide Sequences to the Human and Wheat Proteomes**

Inspection of the PEAKS output indicated that the software truncated, omitted, or otherwise incorrectly parsed the protein accession codes for some peptides (possibly due to the large size of the grain databases that were searched). A Python script was therefore used to generate the “Accession(s)” column reported in the supplemental datasets, ensuring that all protein accession codes corresponding to a given peptide are reported. For downstream data analysis, peptide sequences were assigned to the wheat proteome only if they uniquely mapped to the wheat proteome; sequences that mapped to both the wheat and human proteomes were counted as human peptides. Additionally, peptides mapping to the wheat proteome containing the subsequence YVRPD were assigned as human peptides even if the complete peptide sequence uniquely mapped to the wheat proteome. In preliminary analysis of our proof-of-concept studies presented in **Main Text Figure 2**, we observed that peptides containing this subsequence appeared regardless of gluten dietary status. BLAST searching indicated that this subsequence appears in the immunoglobulin heavy chain joining region. We speculate that the complete peptide sequences were formed as the result of V-(D)-J rearrangement of human immunoglobin genes [10].

**SUPPLEMENTARY FIGURES**

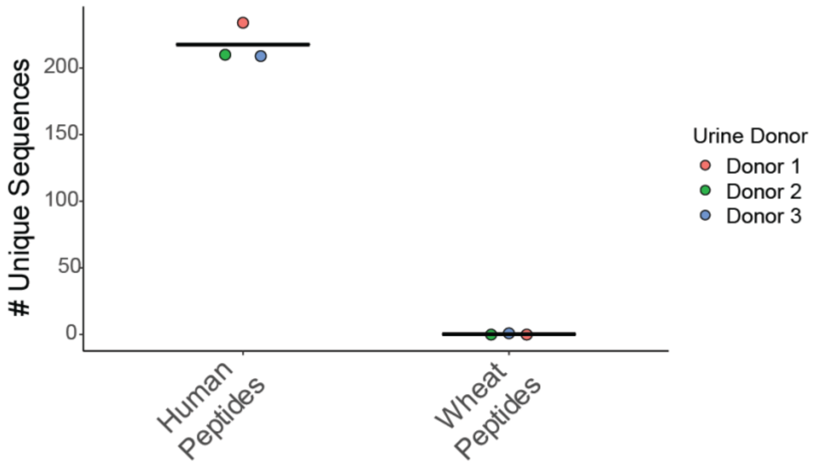


**Supplementary Figure 1**. A standard reversed-phase C18 SPE protocol leads to identification of few human- and wheat-derived urinary peptides. Urine samples (n=3) were processed and analyzed by LC-MS/MS as described in *Pilot Analysis of Urine from Individuals Challenged with Dietary Gluten*. Identified peptide sequences were mapped to the human and wheat proteomes as described in the **Main Text Materials and Methods**. Horizontal lines denote median. Only a single wheat-derived peptide sequence (GQQQPFPPQQPYPQPQPFPS from $\alpha$-gliadin) was detected in the urine of Donor 3. A full list of identified peptides is provided in **Supplementary Dataset 1**.

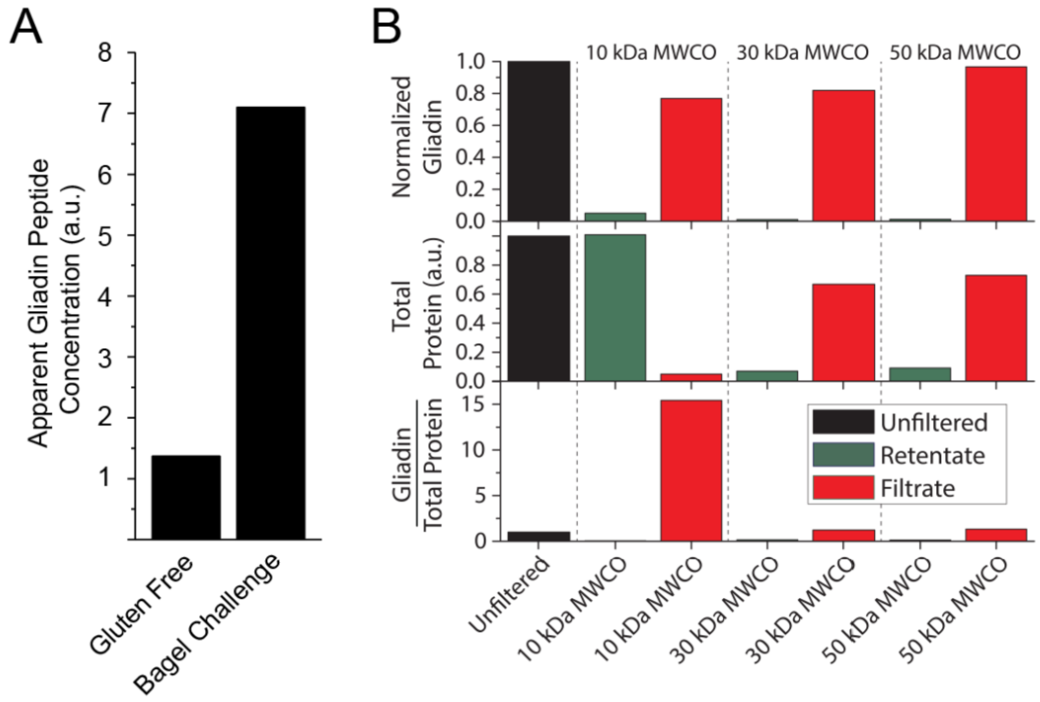


**Supplementary Figure 2.** Identification of a gluten-positive urine reference sample for method optimization to enhance detection of the gluten peptidome. *(A)* Analysis of apparent gliadin peptide content (a.u., arbitrary units) in the urine of an individual before (Gluten Free) and after a challenge with ~18 g dietary gluten (Bagel Challenge), as measured by ELISA using the R5 antibody. An approximately 5-fold higher apparent signal in the R5 ELISA subsequent to the bagel challenge was observed relative to the gluten free sample, where the signal was presumably to due to background binding of the R5 antibody to urinary proteins. This suggested that the "Bagel Challenge" sample was suitable for optimizing our sample extraction and LC-MS/MS method. *(B)* Urinary gliadin peptides can be recovered by ultrafiltration. The "Bagel Challenge" urine sample from *A* was passed through 10, 30, and 50 kDa centrifugal filters, and both the filtrate and retentate were analyzed. The top panel shows the gliadin content of each fraction, as evaluated by R5 ELISA, and normalized by the unfiltered urine sample. The majority of gliadin peptides pass through all ultrafiltration units. The middle panel shows the normalized protein in each fraction, as evaluated by silver-stain densiometry. For 10 kDa filtration, the majority of urinary protein is retained in the retentate. The bottom panel shows the enrichment of gliadin relative to total protein. For both the 30 and 50 kDa membranes, a majority of the R5-reactive gliadin peptides as well as the total protein content were recovered in the filtrate. In contrast, with the 10 kDa filtration device, most of the urinary protein was in the retentate, and a majority of the R5-reactive gliadin peptides passed into the filtrate, resulting in a >15-fold enrichment of gliadin peptides. Accordingly, the 10 kDa MWCO membrane appeared most suitable for efficiently recovering the gluten peptidome while depleting high molecular weight species prior to LC-MS/MS.

A

Sample Processing Workflow ---------------------------------------------------------------->

| | Pre-Filtration Denaturation | Sample Drying | Solid-Phase Extraction | EtOAc Extraction | Sample Drying | Solid-Phase Extraction | Sample Drying | LC Gradient | Orbitrap MS Instrument |
|---|---|---|---|---|---|---|---|---|---|
| **Method A** | | | C18 Column | ✔ | ✔ | | | Short | Elite |
| **Method B** | 20% ACN | ✔ | C18 Column | ✔ | ✔ | | | Short | Elite |
| **Method C** | 2% FA | | C18 Column | ✔ | ✔ | | | Short | Elite |
| **Method D** | 2% FA | | C18 Column | ✔ | ✔ | SCX StageTip | ✔ | Short | Elite |
| **Method E** | 2% FA | | C18 Column | ✔ | ✔ | SCX StageTip | ✔ | Extended | Elite |
| **Method F** | 2% FA | | C18 Column | ✔ | ✔ | SCX StageTip | ✔ | Extended | Fusion |
| **Method G** | 2% FA | | MCX Column | | ✔ | | | Extended | Fusion |

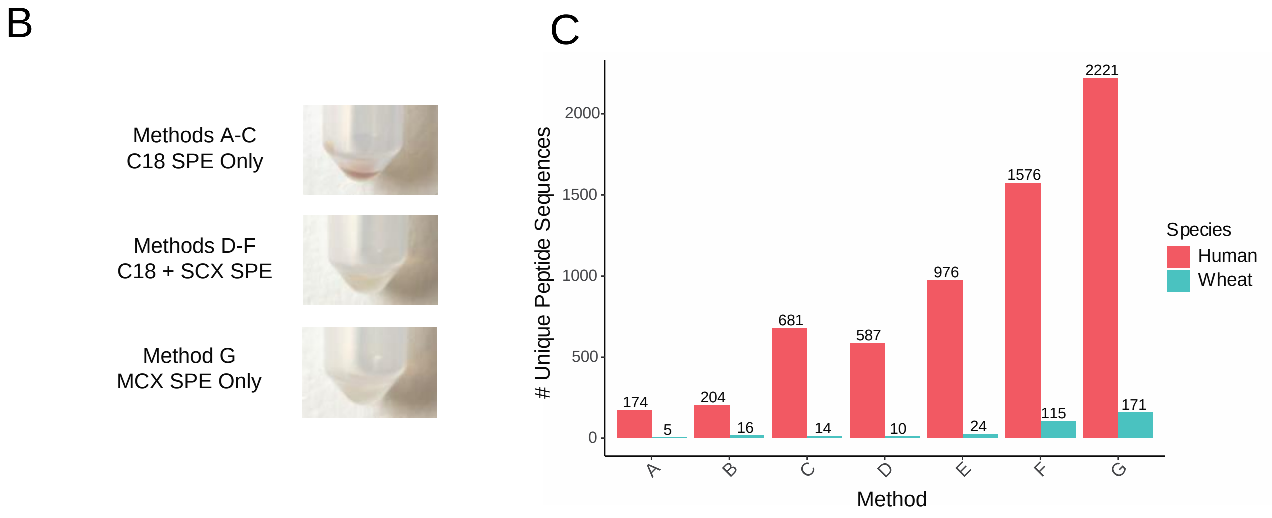


**Supplementary Figure 3.** Optimization of urinary peptidomic cleanup and LC/MS-MS analysis method. *(A)* Key steps in seven different protocols (Methods A-G) that were evaluated. The sample processing workflow moves from left to right, and a particular step was included in a method if the box is highlighted green. *(B)* Representative photos of urinary peptides after processing with the methods shown in *A*. Peptides processed with C18 SPE and ethyl acetate (EtOAc) extraction (Methods A-C) retained a yellow color due to carryover of urinary pigments that were not efficiently removed by SPE or liquid-liquid extraction, consistent with the low solubility of urochrome in ethyl acetate [7]. Addition of an SCX SPE step subsequent to C18 SPE (Methods D-F) appeared to remove the majority of urinary pigments. Urine extracted using a single-step MCX SPE method without ethyl acetate extraction (Method G) was completely colorless. *(C)* Number of human and wheat peptides identified from database searching of LC-MS/MS data after sample processing using Methods A-G. For all methods, 4 mL of the same gluten-positive reference ("Bagel Challenge") urine described in **Supplementary Figure 2** was used. A full list of the peptides identified using each Method is provided in **Supplementary Dataset 2**.

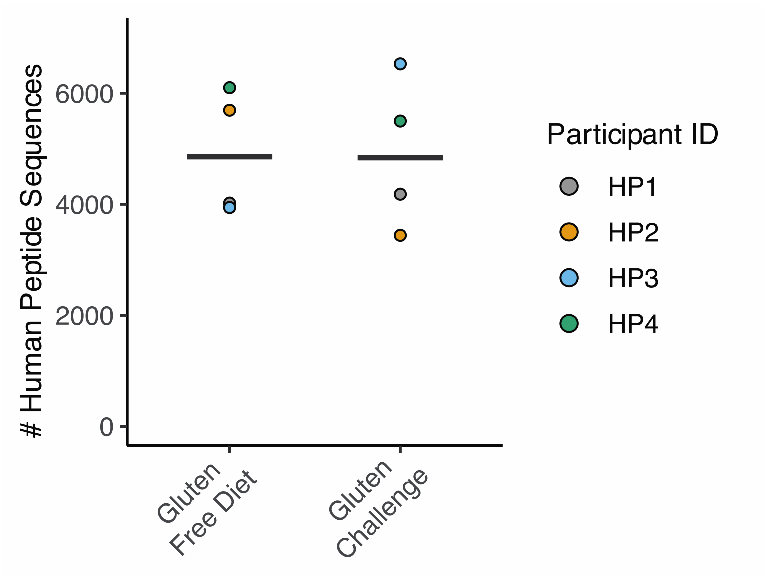


**Supplementary Figure 4.** Number of unique human peptide sequences identified in 8-hour pooled urine collections from 4 healthy participants (HPs) after a 24-hour gluten free diet and after challenge with dietary wheat gluten, as detailed in **Main Text Figure 2a-b**. Horizontal line indicates median. Full details related to the LC-MS/MS identification of these peptide sequences in each participants' urine sample are provided in **Supplementary Dataset 3**.

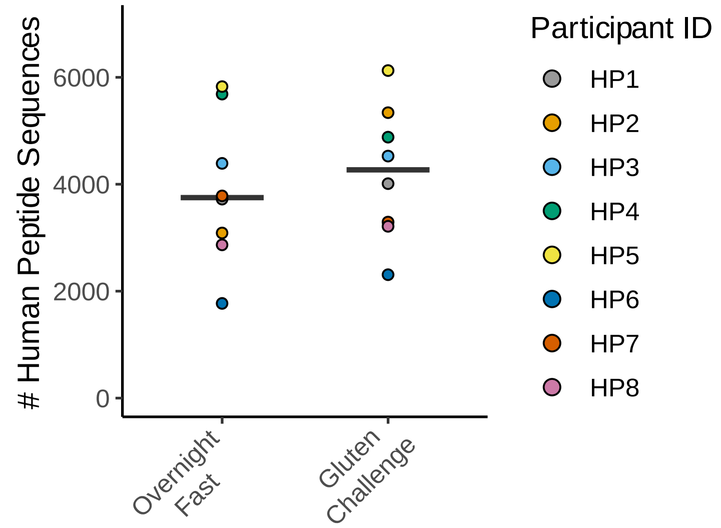


**Supplementary Figure 5.** Number of unique human peptide sequences identified from urine specimens collected from 8 healthy participants (HPs). Single urine specimens were collected after overnight fasting, and an 8-hour pooled urine sample was collected after a dietary gluten challenge, as detailed in **Main Text Figure 2c-d**. Horizontal line indicates median. Full details related to the LC-MS/MS identification of these peptide sequences in each participants' urine sample are provided in **Supplementary Dataset 4**.

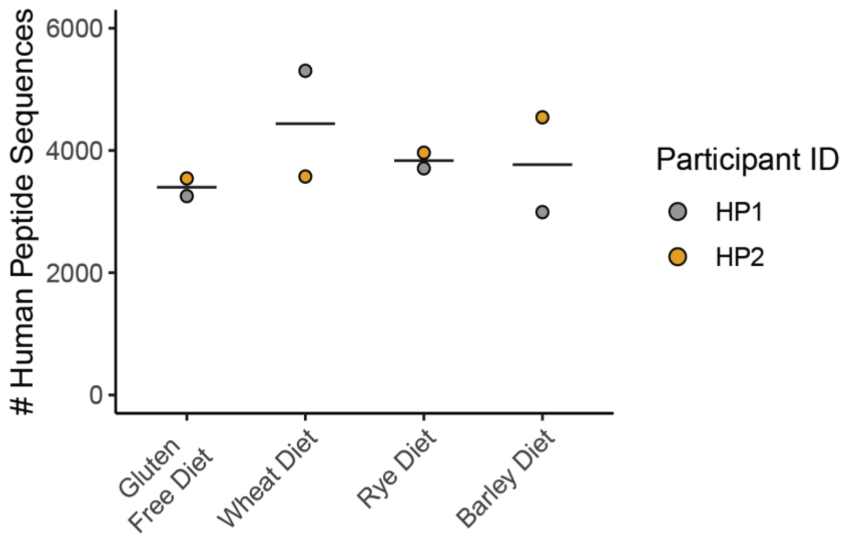


**Supplementary Figure 6.** Number of unique human peptide sequences identified from the urine of 2 healthy participants (HPs) in 8-hour urine collections while maintaining a gluten-free diet, or after dietary challenge with wheat, rye, or barley, as detailed in **Main Text Figure 3**. Horizontal line indicates median. Full details related to the LC-MS/MS identification of these peptide sequences in each participants' urine sample are provided in **Supplementary Dataset 5.**

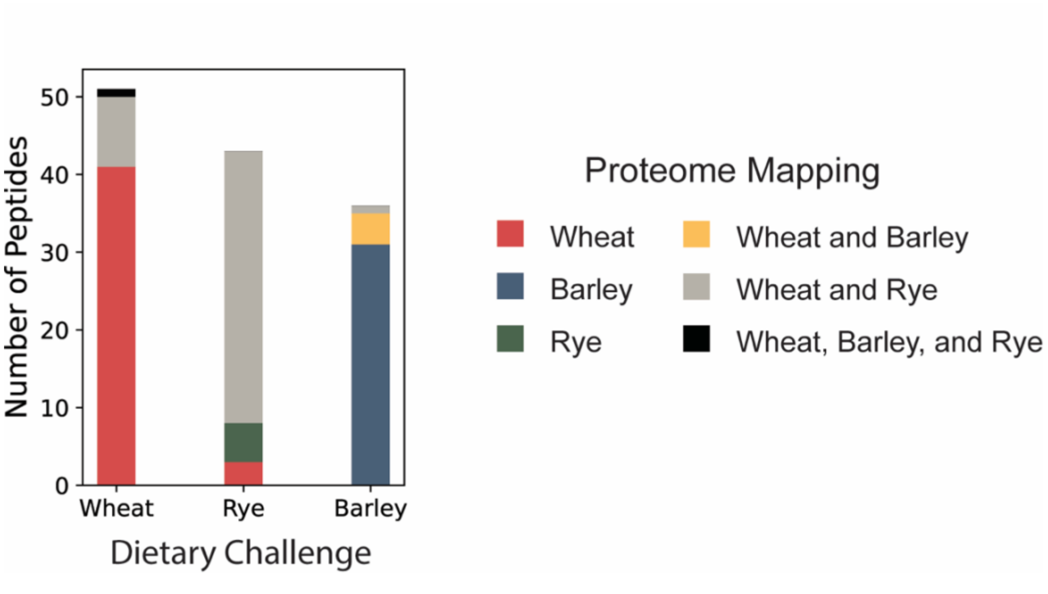


**Supplementary Figure 7.** Peptide-to-proteome mapping of peptides identified from LC-MS/MS analysis of urine from two healthy participants challenged with dietary wheat, rye, and barley as detailed in **Main Text Figure 3.** The large proportion of peptides uniquely mapping to the wheat proteome in the rye dietary challenege is a likely artifact of the small size of the rye proteome currently available in the Uniprot resource, which contains >100x fewer sequences than either the wheat or barley proteomes (**Supplementary Methods**). A full list of observed peptide sequences in each participant and their proteome mappings is provided in **Supplementary Dataset 5.**

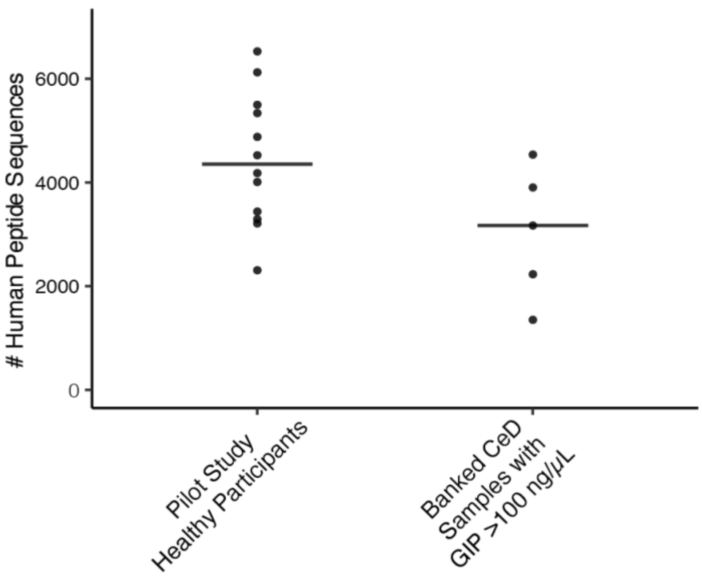


**Supplementary Figure 8.** Comparison of number of unique human peptide sequences identified in 8-hour urine of gluten-challenged healthy participants (pooled data from experiments in **Main Text Figure 2b,d**) and single void urine collections of banked single void CeD urine samples gluten immunogenic peptide (GIP) concentrations of >100 ng/μL (data corresponds to **Main Text Figure 4**).

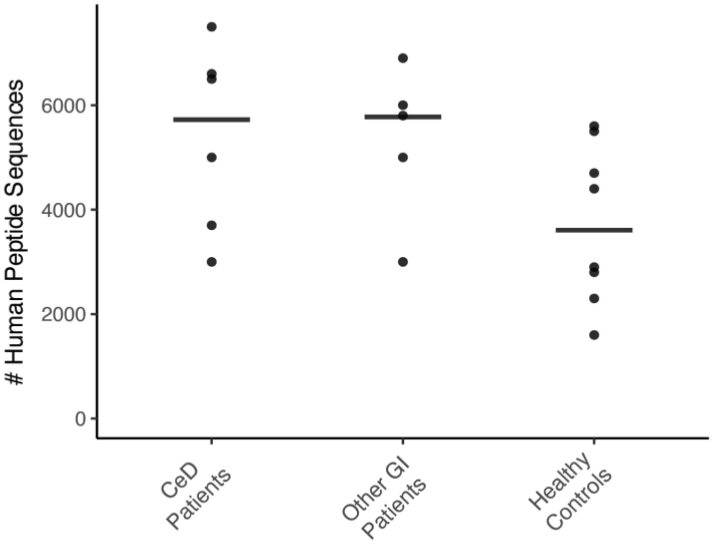


**Supplementary Figure 9.** Number of unique human peptide sequences identified in the urine of 6 patients with CeD, 5 patients with non-celiac gastrointestinal (GI) disorders, and 8 healthy controls in pooled urine samples collected for 8 h subsequent to a dietary challenge with two bagels. Data correspond to **Main Text Figure 5b**. Horizontal line indicates median. Differences between the three groups were not significant ($p > 0.05$, one-way Kruskal-Wallis ANOVA). Full details related to the LC-MS/MS identification of these peptide sequences in each participants' urine sample are provided in **Supplementary Dataset 7.**

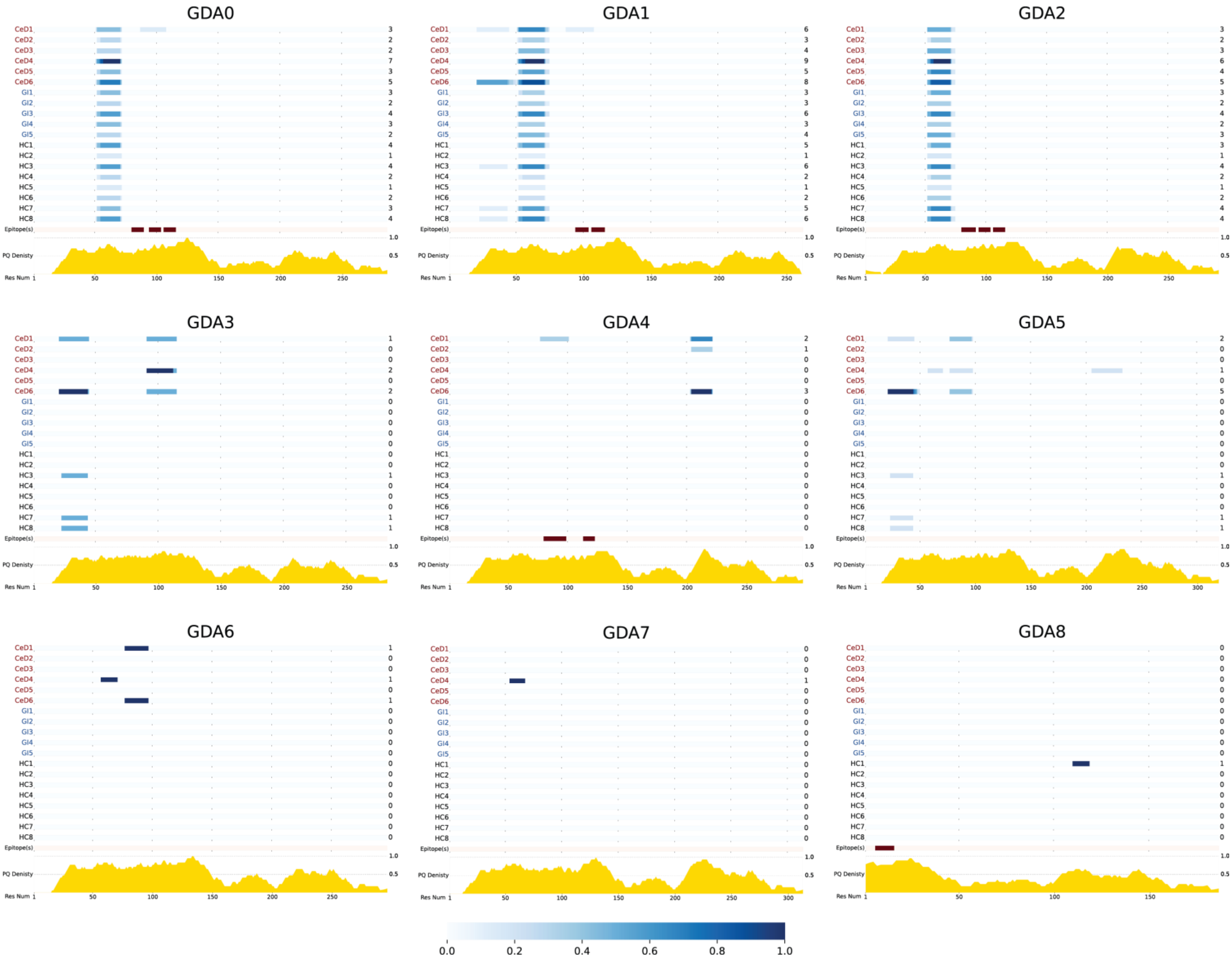
GDA0
GDA1
GDA2
GDA3
GDA4
GDA5
GDA6
GDA7
GDA8
CeD1
CeD2
CeD3
CeD4
CeD5
CeD6
GI1
GI2
GI3
GI4
GI5
HC1
HC2
HC3
HC4
HC5
HC6
HC7
HC8
Epitope(s)
PQ Denisty
Res Num
1.0
0.5
1
50
100
150
200
250
300
3 2 2 7 3 5 3 2 4 3 2 4 1 4 2 1 2 3 4
6 3 4 9 5 8 3 3 6 3 4 5 1 6 2 1 2 5 6
3 2 3 6 4 5 3 2 4 2 3 3 1 4 2 1 2 4 4
1 0 0 2 0 2 0 0 0 0 0 0 0 1 0 0 0 1 1
2 1 0 0 0 3 0 0 0 0 0 0 0 0 0 0 0 0 0
2 0 0 1 0 5 0 0 0 0 0 0 0 1 0 0 0 1 1
1 0 0 1 0 1 0 0 0 0 0 0 0 0 0 0 0 0 0
0 0 0 1 0 0 0 0 0 0 0 0 0 0 0 0 0 0 0
0 0 0 0 0 0 0 0 0 0 0 1 0 0 0 0 0 0 0
0.0
0.2
0.4
0.6
0.8
1.0

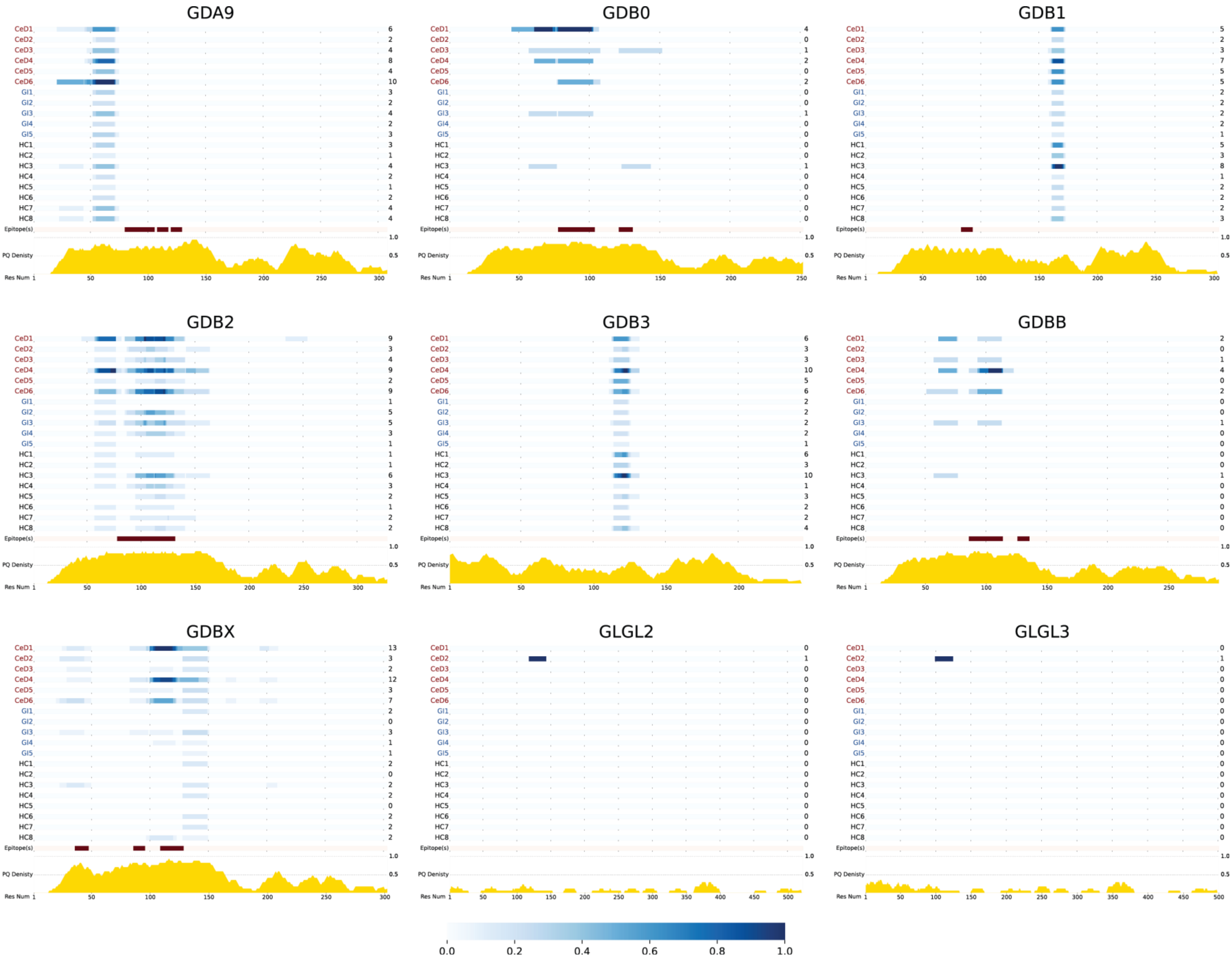
GDA9
GDB0
GDB1
GDB2
GDB3
GDBB
GDBX
GLGL2
GLGL3
CeD1
CeD2
CeD3
CeD4
CeD5
CeD6
GI1
GI2
GI3
GI4
GI5
HC1
HC2
HC3
HC4
HC5
HC6
HC7
HC8
Epitope(s)
PQ Denisty
Res Num
GDA9: 6 2 4 8 4 10 3 2 4 2 3 3 1 4 2 1 2 4 4
GDB0: 4 0 1 2 0 2 0 0 1 0 0 0 0 1 0 0 0 0 0
GDB1: 5 2 3 7 5 5 2 2 2 2 1 5 3 8 1 2 2 2 3
GDB2: 9 3 4 9 2 9 1 5 5 3 1 1 1 6 3 2 1 2 2
GDB3: 6 3 3 10 5 6 2 2 2 2 1 6 3 10 1 3 2 2 4
GDBB: 2 0 1 4 0 2 0 0 1 0 0 0 0 1 0 0 0 0 0
GDBX: 13 3 2 12 3 7 2 0 3 1 1 2 0 2 2 0 2 2 2
GLGL2: 0 1 0 0 0 0 0 0 0 0 0 0 0 0 0 0 0 0 0
GLGL3: 0 1 0 0 0 0 0 0 0 0 0 0 0 0 0 0 0 0 0
1.0
0.5
0.0 0.2 0.4 0.6 0.8 1.0

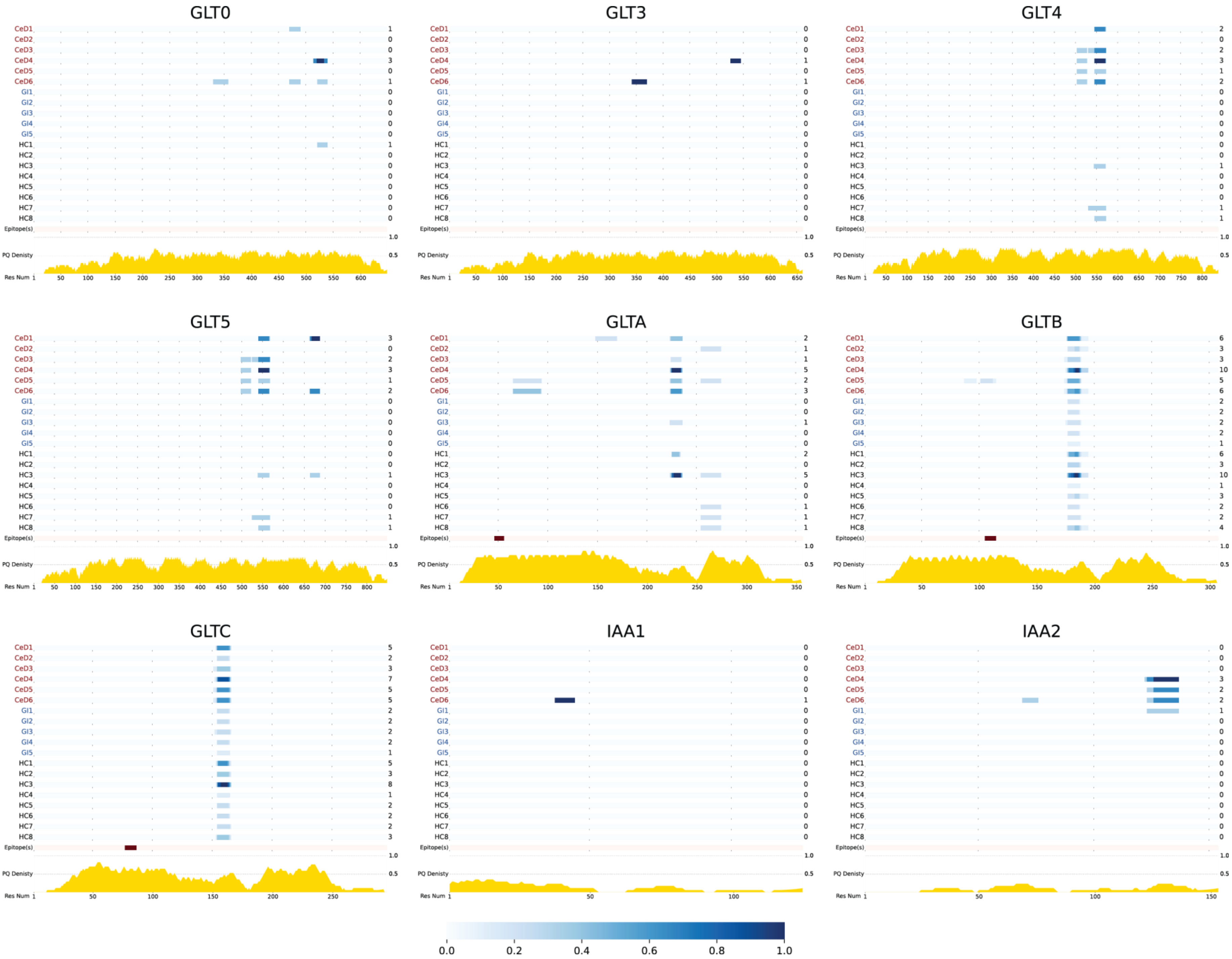
GLT0
GLT3
GLT4
GLT5
GLTA
GLTB
GLTC
IAA1
IAA2
CeD1
CeD2
CeD3
CeD4
CeD5
CeD6
GI1
GI2
GI3
GI4
GI5
HC1
HC2
HC3
HC4
HC5
HC6
HC7
HC8
Epitope(s)
PQ Denisty
Res Num
0.0
0.2
0.4
0.6
0.8
1.0

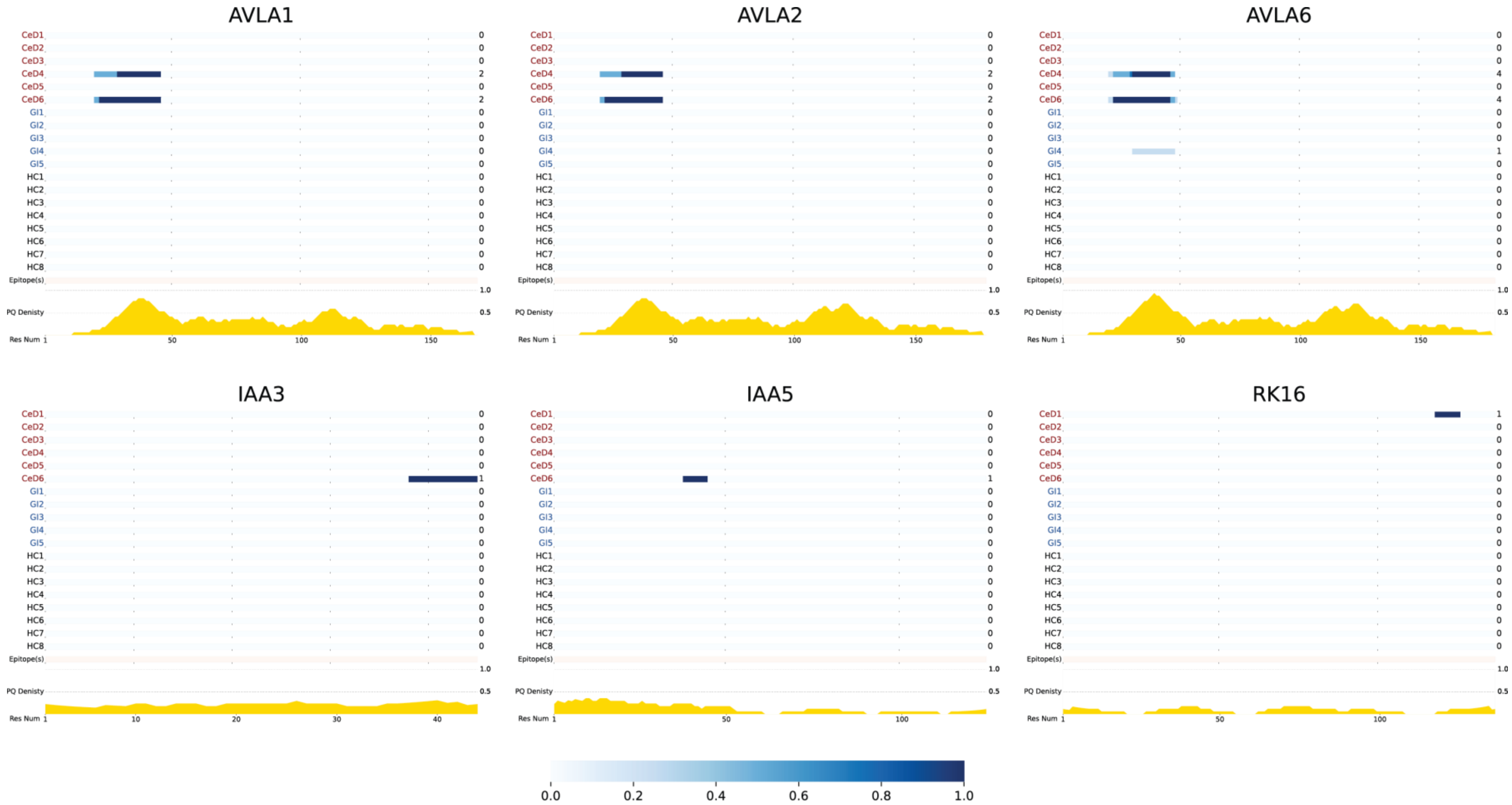


**Supplementary Figure 10.** Heatmap representation highlighting the locations in the wheat proteome covered by urinary peptides in each clinical study participant. Detected peptide sequences were mapped to the wheat proteome through exact sequence alignment, as shown in **Main Text Figure 5g.** For this mapping, only proteins in the wheat SwissProt proteome were considered. If a peptide's sequence aligned to multiple protein sequences or multiple sites within a single protein, then all of its potential locations in the proteome are indicated in this heatmap. Each row of the heatmap reflects the frequency at which the corresponding protein region was identified in each study participant's urine (CeD: patient with celiac disease; GI: patient with non-celiac gastrointestinal disorder; HC: healthy control), normalized to the maximum frequency in all participants. The frequencies shown are based on a simple count of unique peptides and are not adjusted for multiple spectrum identifications of the same peptide or intensity-based peptide abundance inferences. Below the heatmaps, regions of the proteome that contain known CeD-relevant T-cell epitopes are indicated in dark red[11], and the density of Pro and Gln residues (PQ density) is plotted in yellow. The density value at a given position is calculated as the fraction of residues in a window of 17 (residue at the current position, 8 proceeding residues, and 8 succeeding residues) that contains a proline or glutamine. For the first and last 8 positions of a protein sequence, preceding and succeeding window sizes are limited, respectively.

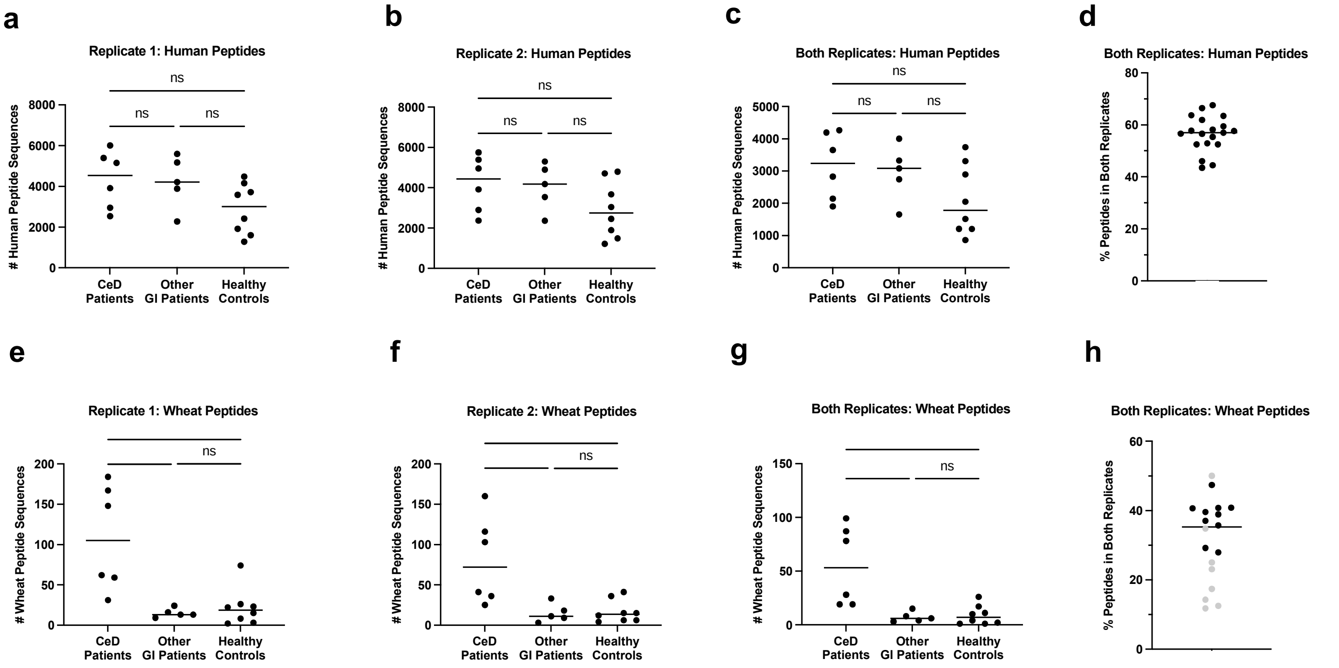


**Supplementary Figure 11** (reproduced here for convenience)*.* Technical reproducibility of peptide identifications. Urine samples from CeD patients (n=6), patients with non-CeD gastrointestinal disorders (n=5) or healthy controls (n=8) were independently prepared on separate days and analyzed in independent LC-MS/MS runs. (a) Human peptide sequences identified in Replicate 1. (b) Human peptide sequences identified in Replicate 2. (c) Human peptide sequences identified only in both replicates. (d) Human peptide sequences identified only in both replicates expressed as a percentage of total peptides. (e) Wheat peptide sequences identified in Replicate 1. (f) Wheat peptide sequences identified in Replicate 2. (g) Wheat peptide sequences identified only in both replicates. (h) Wheat peptide sequences identified only in both replicates expressed as a percentage of total peptides. Samples with fewer than 20 wheat peptides identified in either replicate are denoted as gray circles. *$p < 0.05$; **$p < 0.01$. One-way Kruskal-Wallis ANOVA/Dunn's multiple comparison test.

#### SUPPLEMENTARY TABLES

**Supplementary Table 1.** Summary of peptides containing CeD-relevant T-cell epitopes identified in the urine of 8 healthy volunteers (from all peptide sequences listed in **Supplementary Datasets 3 and 4).**

| **Epitope Name(s)[a]** | **Epitope Sequence[a]** | **# Unique Peptides** | **Peptide Sequences with/ Epitope[b]** |
|---|---|---|---|
| DQ2.5-glia-g1<br>/DQ8.5-glia-g1/<br>DQ8-glia-g2 | PQQSFPQQQ | 1 | PQQQFPQPQQ**PQQSFPQQQ**QPLIQPYLQQQMNPC(+57.02)KNYLLQQC(+57.02)NP |
| DQ2.5-glia-g3/<br>DQ8-glia-g1b | QQPQQPYPQ | 1 | FLQPQQPFP**QQPQQPYPQ**QPQQPFPQ |
| DQ2.5-glia-g4b | PQPQQQFPQ | 4 | SQQPQQPF**PQPQQQFPQ**PQQPQ,<br>SQQPQQPF**PQPQQQFPQ**PQQPQQ,<br>SQQPQQPF**PQPQQQFPQ**PQQP,<br>Q(-17.03)QPHQPF**PQPQQQFPQ**PQQPQQS |
| DQ2.5-glia-g4c/<br>DQ8-glia-g1a | QQPQQPFPQ | 33 | EQTIS**QQPQQPFPQ**QPHQPQQPYPQQQPYGSSL,<br>SQQPEQTIS**QQPQQPFPQ**QPHQPQQPYPQQQPYGSSL,<br>S**QQPQQPFPQ**QPHQPQQPYPQQ,<br>S**QQPQQPFPQ**PQQQFPQPQQPQ,<br>S**QQPQQPFPQ**PQQQFPQPQQPQQ,<br>S**QQPQQPFPQ**PQQQFPQPQQP,<br>S**QQPQQPFPQ**QPHQPQQPYPQQQP,<br>S**QQPQQPFPQ**QPHQPQQPYPQ,<br>PQLPFP**QQPQQPFPQ**PQQP,<br>QPQQPFPQT**QQPQQPFPQ**QPQQPFPQ,<br>PQQPQLPFP**QQPQQPFPQ**PQ,<br>TQQPQQPFP**QQPQQPFPQ**T,<br>PLQPQQPFP**QQPQQPFPQ**P,<br>PLQPQQPFP**QQPQQPFPQ**PQ,<br>PLQPQQPFP**QQPQQPFPQ**QP,<br>TQQPQQPFP**QQPQQPFPQ**QPQQPFP,<br>AQLPFPQQP**QQPFPQPQQ**PQQPFPQ,<br>S**QQPQQPFPQ**QPHQPQQPYP,<br>T**QQPQQPFPQ**QPQQPFP**QQPQQPFPQ**,<br>**QQPQQPFPQ**QPHQPQQPYP,<br>LQQPQQPLPQP**QQPQQPFPQ**QQQPL,<br>QPQQPFPQT**QQPQQPFPQ**QPQQPFP,<br>T**QQPQQPFPQ**QPQQPFPQT**QQPQQPFPQ**,<br>LQPQQPFP**QQPQQPFPQ**T**QQPQQPFPQ**,<br>FLQPQQPFPQQPQQPYP**QQPQQPFPQ**,<br>PQQPQLPFP**QQPQQPFPQ**P**QQPQQPFPQ**,<br>PQQPQLPFP**QQPQQPFPQ**PQQPQQ,<br>IS**QQPQQPFPQ**QPHQPQQPYPQQQPYGSSL,<br>SQQLEQTIS**QQPQQPFPQ**QPHQPQQPYPQQQPYGSSL,<br>PLQPQQPFP**QQPQQPFPQ**PQLPFPQQSE,<br>PLQPQQPFP**QQPQQPFPQ**PQLPFPQQ<br>ESQQPFP**QQPQQPFPQ**PQ, |
| DQ2.5-glia-g4e | LQPQQPFPQ | 6 | P**LQPQQPFPQ**QPQQPFPQQP,<br>**LQPQQPFPQ**QPQQPFPQTQQPQQPFPQ,<br>F**LQPQQPFPQ**QPQQPYPQQPQQPFPQ,<br>P**LQPQQPFPQ**QPQQPFPQPQLPFPQQSE,<br>P**LQPQQPFPQ**QPQQPFPQPQLPFPQQ,<br>P**LQPQQPFPQ**QPQQPFPQPQ |
| DQ2.5-glia-g5 | QQPFPQQPQ | 14 | QPQQPFPQT**QQPQQPFPQ**QPQQPFPQ,<br>T**QQPQQPFPQ**QPQQPFPQT,<br>PLQPQQPFP**QQPQQPFPQ**P,<br>PLQPQQPFP**QQPQQPFPQ**PQ,<br>PLQPQQPFP**QQPQQPFPQ**QP,<br>T**QQPQQPFPQ**QPQQPFPQQPQQPFP,<br>T**QQPQQPFPQ**QPQQPFP**QQPQQPFPQ**,<br>QPQQPFPQT**QQPQQPFPQ**QPQQPFP,<br>T**QQPQQPFPQ**QPQQPFPQT**QQPQQPFPQ**,<br>LQPQQPFP**QQPQQPFPQ**T**QQPQQPFPQ**,<br>FLQPQQPFPQQPQQPYP**QQPQQPFPQ**,<br>PLQPQQPFP**QQPQQPFPQ**PQLPFPQQSE,<br>PLQPQQPFP**QQPQQPFPQ**PQLPFPQQ,<br>ESQQPFP**QQPQQPFPQ**PQ |

| | | | |
|---|---|---|---|
| DQ2.5-glut-L1/<br>DQ2.2-glut-L1 | PFSQQQPV | 6 | PQQPPFSQQQLPPFSQQLP**PFSQQQQPV**,<br>PQQPPFSQQQQQQQQQQQP**PFSQQQQPV**L,<br>PQQPPFSQQQQQQQQQQP**PFSQQQQPV**L,<br>PQQPPFSQQQQQQQQQQP**PFSQQQQPV**,<br>PQQPPFSQQQQQPILPQQP**PFSQQQQPV**,<br>PQQPPFSQQQQQPILPQQP**PFSQQQQPV**L, |

[a]Epitope nomenclature and sequences are defined in ref. 11.
[b](+57.02) denotes cysteine alkylation by iodoacetamide during sample workup.

**Supplementary Table 2.** Summary of peptides containing CeD-relevant T-cell epitopes found from the clinical study presented in **Main Text Figure 5** that were detected exclusively in patients with CeD.

| **Epitope Name[a]** | **Epitope Sequence[a]** | **# Unique Peptides** | **Peptide Sequences w/ Epitope[b]** |
|---|---|---|---|
| DQ2.5-glia-α1a | PFPQPQLPY | 2 | Q(-17.03)LQ**PFPQPQLPY**PQPHLPYPQPQP,<br>QLQ**PFPQPQLPY**PQPQLPYPQPQP,<br>Q(-17.03)LQ**PFPQPQLPY**PQPHLPYPQPQP, |
| DQ2.5-glia-α1b | PYPQPQLPY | 1 | QLQPFPQPQL**PYPQPQLPY**PQPQP, |
| DQ2.5-glia-α2 | PQPQLPYPQ | 2 | Q(-17.03)LQPF**PQPQLPYPQ**PHLPYPQPQP,<br>QLQPF**PQPQLPYPQ**PQLPYPQPQP,<br>Q(-17.03)LQPF**PQPQLPYPQ**PHLPYPQPQP |
| DQ2.5-glia-α3 | FRPQQPYPQ | 1 | PYSQPQP**FRPQQPYPQ**PQPQY,<br>PYSQPQP**FRPQQPYPQ**PQPQY, |
| DQ2.5-glia-g4α | SQPQQQFPQ | 1 | **SQPQQQFPQ**PQQPQQSFPQQQPPFIQPSLQQ |
| DQ2.5-glia-ω2 | PQPQQPFPW | 1 | QQPQQPF**PQPQQPFPW**QPQQPFP |
| DQ2.5-glut-L2 | FSQQQQSPF | 1 | Q(-17.03)ISQQQQQPPFSQQQQPP**FSQQQQSPF**SQQQQQPPFL |

[a]Epitope nomenclature and sequences are defined in ref.11.
[b](+57.02) denotes cysteine alkylation by iodoacetamide during sample workup. (-17.03) denotes formation of pyroglutamic acid from glutamine.

**Supplementary Table 3.** Clinical information for CeD study participants.

| ID | Age | Sex | IgA Anti-TG2 | IgA Anti-DGP | HLA Status | Endoscopic Findings | Diagnosis |
|---|---|---|---|---|---|---|---|
| **GI1** | 30 | F | <1.0 | 1.2 | HLA DQ2 +/+ | No villous abnormality or increased intraepithelial lymphocytosis | Non-CeD GI |
| **CeD1** | 61 | F | 22 | >140.0 | ND | Mild villous atrophy and increased intraepithelial lymphocytosis | CeD |
| **GI2** | 45 | F | <1 | 4 | ND | No villous abnormality or increased intraepithelial lymphocytosis | Non-CeD GI |
| **CeD2** | 20 | F | 52 | 1.4 | ND | Villous blunting and increased intraepithelial lymphocytosis | CeD |
| **CeD3** | 34 | F | >128.0 | 73 | ND | Villous blunting and increased intraepithelial lymphocytosis | CeD |
| **GI3** | 60 | M | <1.0 | ND | HLA DQ8 +/+ | No villous abnormality or increased intraepithelial lymphocytosis | Non-CeD GI |
| **GI4** | 20 | F | ND | 2.2 | HLA DQ2 and DQ8 -/- | Not performed due to negative HLA typing | Non-CeD GI |
| **CeD4** | 31 | F | 82 | ND | ND | Villous blunting and increased intraepithelial lymphocytosis | CeD |
| **CeD5** | 37 | F | 47.6 | 54 | HLA DQ2 +/? | No villous atrophy but increased intraepithelial lymphocytosis | CeD |
| **CeD6** | 32 | F | ND | >140 | ND | Severe blunting and increased intraepithelial lymphocytosis | CeD |
| **GI5** | 61 | F | 3.1 | <1.0 | HLA DQ8 +/- | Patient refused endoscopy based on negative serology | Non-CeD GI |

**Supplementary Datasets 1-7** contain full information related to the identification of peptide sequences in each study participant's urine sample(s), including PEAKS software identification scores (-10lgP), detected mass-to-charge ratios (m/z), LC retention times (RT, given in minutes), and experimental mass errors (ppm compared to the theoretical m/z). Within each dataset, sheets ending in "All" contain all peptide sequences (human and wheat, and where applicable, barley and rye). Sheets ending in "Wheat" or "Grain" have those sequences that map uniquely to the wheat, and where applicable, rye and barley proteomes. These sheets containing only grain peptides have additional columns titled "Manually Corrected Peptide Sequence" and "Included in Final Dataset". These columns respectively indicate where peptide sequences were reassigned from deamidated to native or excluded entirely from analysis as described above in **Validation of Wheat-Derived Peptide Sequences Identified by PEAKS Software**.

**SI REFERENCES**

_


1. Rogers, J. C. & Bomgarden, R. D. Sample preparation for mass spectrometry-based proteomics; from proteomes to peptides. *Adv. Exp. Med. Biol.* **919**, 43-62 (2016).
2. Boucher, M. J. et al. Integrative proteomics and bioinformatic prediction enable a high-confidence apicoplast proteome in malaria parasites. *PLoS Biol* **16**, e2005895 (2018).
3. Smith, C. R. et al. Deciphering the peptidome of urine from ovarian cancer patients and healthy controls. *Clin. Proteomics* **11**, 23 (2014).
4. Sigdel, T. K. et al. Optimization for peptide sample preparation for urine peptidomics. *Clin. Proteomics* **11**, 7 (2014).
5. Di Meo, A. et al. An integrated proteomic and peptidomic assessment of the normal human urinome. *Clin. Chem. Lab. Med.* **55**, 237-247 (2017).
6. Van, J. A. D. et al. Peptidomic analysis of urine from youths with early type 1 diabetes reveals novel bioactivity of uromodulin peptides in vitro. *Mol. Cell Proteomics* **19**, 501-517 (2020).
7. Drabkin, D. L. The normal pigment of the urine. *J. Biol. Chem* **75**, 481 (1927).
8. Rappsilber, J., Mann, M. & Ishihama, Y. Protocol for micro-purification, enrichment, pre-fractionation and storage of peptides for proteomics using StageTips. *Nat. Protoc.* **2**, 1896-1906 (2007).
9. Nepomuceno, A. I., Gibson, R. J., Randall, S. M. & Muddiman, D. C. Accurate identification of deamidated peptides in global proteomics using a quadrupole orbitrap mass spectrometer. *J. Proteome Res.* **13**, 777-785 (2014).
10. Watson, C. T. & Breden, F. The immunoglobulin heavy chain locus: genetic variation, missing data, and implications for human disease. *Genes Immun.* **13**, 363-373 (2012).
11. Sollid, L. M. et al. Update 2020: nomenclature and listing of celiac disease-relevant gluten epitopes recognized by $CD4^+$ T cells. *Immunogenetics* **72**, 85-88 (2020).
